# AMFR provides an ERAD bypass mechanism to maintain proteostasis under canonical E3 ligase deficiency

**DOI:** 10.1101/2025.05.07.652780

**Authors:** Uta Nakayamada, Aika Fujii, Saori Kato, Yuka Kamada, Tsukasa Okiyoneda

**Affiliations:** Department of Biomedical Sciences, School of Biological and Environmental Sciences, Kwansei Gakuin University, Sanda 669-1330, Hyogo, Japan

**Keywords:** ERAD, ubiquitin ligase, E3, RNF5, RNF185, AMFR, CFTR

## Abstract

Abnormal proteins in the endoplasmic reticulum (ER) are eliminated via distinct ER-associated degradation (ERAD) pathways, each regulated by specific E3 ubiquitin ligases. While these pathways seem to work cooperatively to maintain ER proteostasis, their individual roles and potential compensatory mechanisms remain poorly defined in mammalian cells. In this study, we utilized multiple E3 ligase knockouts/knockdowns combined with a highly sensitive HiBiT-based ERAD assay to investigate pathway complementarity. We discovered that in the absence of RNF5/185 function, AMFR—a Hrd1 ortholog primarily involved in the ERAD-M branch—could partially compensate by facilitating degradation of mutant CFTR. In contrast, SYVN1, another Hrd1 ortholog, failed to show a similar effect. These findings reveal a novel bypass mechanism mediated by AMFR and demonstrate the functional flexibility of the ERAD network. Our results provide new insight into how E3 ligases maintain proteostasis under compromised conditions, advancing our understanding of ERAD pathway coordination in mammalian systems.

## Introduction

In the endoplasmic reticulum (ER), aberrant proteins, including misfolded and unassembled proteins, are selectively recognized by ER quality control (ERQC) mechanisms and primarily degraded through the ubiquitin (Ub)-proteasome system.^1,2^ In coordination with molecular chaperones and other accessory factors, Ub ligases facilitate ER-associated degradation (ERAD), a process in which defective proteins are retrotranslocated from the ER to the cytoplasm for degradation by the proteasome.^1–4^ The specific ERAD pathway utilized depends on the localization of conformational defects, with different Ub ligases directing proteins to distinct ERAD branches.^2,5^ The yeast Saccharomyces cerevisiae possesses three distinct ERAD branches, each defined by a specific Ub ligase complex: Hrd1, Doa10, and Asi.^2,5–8^ These ERAD pathways exhibit substrate specificity and are localized to different regions of the ER, collectively targeting a wide range of misfolded proteins for degradation. The Hrd1 complex plays a unique role in eliminating soluble, ER luminal proteins classified as ERAD-L substrates. Additionally, it participates in the degradation of ERAD-M substrates with structural defects in the membrane.^9^ The Doa10 complex is primarily responsible for degrading ERAD-C substrates, recognizing misfolded cytosolic domains on certain membrane proteins.^10^ Furthermore, Doa10 can detect degradation signals within the ER membrane, facilitating the degradation of ERAD-M substrates, such as unassembled subunits of ER membrane complexes and mislocalized membrane proteins.^11,12^ The Asi complex, uniquely localized to the inner nuclear membrane (INM)—a specialized ER subdomain exposed to the nucleoplasm—is specifically involved in the degradation of mislocalized proteins within the INM.^13^

In mammals, the E3 ligases AMFR (Gp78), Synoviolin 1 (SYVN1), and TEB4 (MARCHF6) share a similar domain and topological organization with their yeast counterparts, but exhibit limited sequence homology. Among them, AMFR and SYVN1 are orthologous to yeast Hrd1, whereas MARCHF6 is the orthologue of yeast Doa10.^2,8,14^ Similar to yeast, SYVN1 in mammalian cells plays a crucial role in both the ERAD-L and ERAD-M pathways, facilitating the degradation of aberrant proteins with conformational defects in the ER lumen and intramembrane regions.^15–17^ AMFR and MARCHF6 are involved in ERAD-M and ERAD-C pathways, respectively.^18^ The classification of ERAD-M substrates can be complex, as missense mutations or assembly defects in membrane proteins may either disrupt or promote their incorporation into the lipid bilayer.^19^ These mutations may also induce major structural changes, particularly at domain interfaces. In contrast, ERAD-C substrates include large multipass transmembrane proteins with cytosolic domain mutations, such as the ΔF508 mutant of cystic fibrosis transmembrane conductance regulator (CFTR),^20^ improperly integrated tail-anchored proteins,^21^ and cytoplasmic proteins with exposed hydrophobic regions at domain or subunit interfaces.^22^ In addition to the previously mentioned E3 Ub ligases, several ER membrane-bound Ub ligases have been implicated in ERAD in mammalian cells, including RNF5 (RMA1), RNF185, RNF139 (TRC8), RNF145, RNF170, and TMEM129.^2,8^ Although these distinct ERAD branches are generally regarded as functionally independent, the large number of ER-associated Ub ligases complicates the evaluation of their functional redundancy. Moreover, the degree to which these pathways can compensate for each other when specific ERAD branches are impaired remains largely unclear.

The multispanning membrane protein CFTR is a cAMP-regulated chloride (Cl⁻) channel located at the plasma membrane (PM), and its mutations cause cystic fibrosis (CF), one of the most common genetic disorders.^23^ CFTR consists of two membrane-spanning domains (MSD1 and MSD2), two large cytosolic nucleotide-binding domains (NBD1 and NBD2), and a regulatory (R) domain.^23^ The most common CF-associated mutant, ΔF508-CFTR, results from the deletion of a phenylalanine at position 508 in NBD1, causing misfolding in the ER. This misfolded protein undergoes ubiquitination and is subsequently degraded through the ERAD pathway.^24^ ΔF508-CFTR has been classified as an ERAD-C substrate due to its mutation in the cytosolic NBD1.^25,26^ Multiple Ub E3 ligases have been identified as key regulators of CFTR ERAD. The ER-embedded Ub ligase RNF5 and its paralog RNF185 are thought to mediate the primary CFTR ERAD pathway.^27–29^ Additionally, cytosolic HERC3 and the chaperone-associated ligase CHIP (STUB1) contribute to alternative ERAD pathways.^29,30^ AMFR has been reported to function as an E4-like enzyme downstream of the RNF5-mediated ERAD branch, further facilitating CFTR degradation.^31^ Despite the dominant role of RNF5 and RNF185 (RNF5/185) in CFTR degradation, previous studies have shown that some ΔF508-CFTR is still eliminated in their absence.^29,32^ This suggests that when the primary ERAD pathway is impaired, alternative compensatory degradation mechanisms, referred to as "ERAD bypass", may help maintain ER proteostasis by eliminating misfolded membrane proteins. However, the molecular mechanisms driving ERAD bypass, which likely involves redundant E3 ligases, remain largely unexplored.

In this study, quantitative ERAD analysis using luminescent tags and multiple knockdown/knockout analyses uncovered a compensatory role of AMFR when the primary ERAD branch for membrane proteins is impaired. The findings propose a mechanism in which ERAD branches complement each other by providing an ERAD bypass pathway, ensuring the degradation of misfolded proteins at the ER, with AMFR playing a crucial role in this process.

## Results

### AMFR KD markedly increases ΔF508-CFTR levels in RNF5/185 DKO cells

Our previous study, utilizing a HiBiT-based quantitative ERAD assay, demonstrated that ΔF508-CFTR is primarily degraded through the RNF5/185-dependent ERAD pathway. ^29^ We hypothesized that in RNF5/185 double knockout (DKO) cells, compensatory ERAD mechanisms may be upregulated to eliminate aberrant proteins that are normally targeted by RNF5/185. To identify candidate genes that may compensate for RNF5/185 function, we compared the mRNA expression levels of ERQC-associated Ub E3 ligases involved in ERAD branches in mammalian cells and other ERAD-related components ^2,8^ between wild-type (WT) and RNF5/185 DKO cells. RNA sequencing (RNA-seq) analysis revealed that several ERAD-related genes were upregulated at the mRNA level (Fig. 1A). To determine whether these upregulated genes influence ERAD activity, we assessed the impact of their knockdown (KD) on CFTR PM expression, as ERAD inhibition may improve ΔF508-CFTR PM levels. Among the upregulated candidates including Ub E3 ligases involved in ERAD branches,^2,8^ only KD of AMFR or UBR4 significantly increased the PM expression of ΔF508-CFTR-Nluc(Ex) in RNF5/185 DKO cells ^33^ (Fig. 1B). In contrast, KD of AMFR or UBR4 had no significant effect on the PM levels of ΔF508-CFTR-Nluc(Ex) in WT cells (Fig. S1). Further Western blot analysis revealed that AMFR KD, but not UBR4 KD, led to a substantial increase in both the immature and mature forms of ΔF508-CFTR in RNF5/185 DKO cells (Fig. 1C). Based on these findings, we focused on AMFR for further investigation.

**Figure 1.**
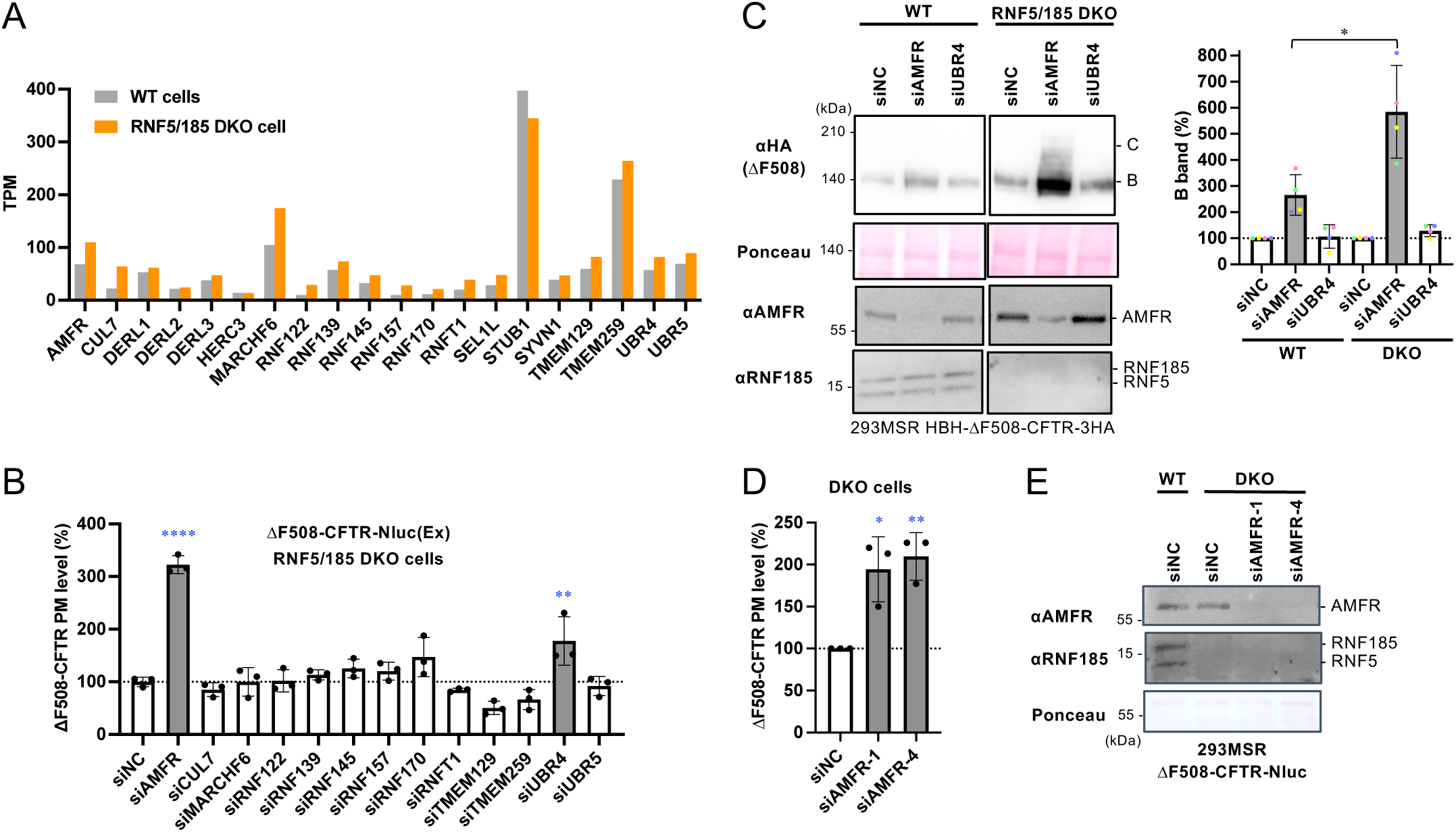
Identification of AMFR as an E3 ligase limiting ΔF508-CFTR expression in RNF5/185 DKO cells. (A) RNAseq analysis highlighting ERAD-related components, including ER-localized E3 ligases, in 293MSR WT and RNF5/185 DKO cells, with mRNA expression shown as transcripts per million (TPM)s(n=1). (B) siRNA screening of E3 ligases limiting cell surface ΔF508-CFTR-Nluc(Ex) in RNF5/185 DKO cells, following transfection with 10 nM siNC (negative control) or pooled siRNA (n=3). (C) Western blot analysis of HBH-ΔF508-CFTR-3HA in 293MSR WT and RNF5/185 DKO cells transfected with 10 nM pooled siRNA, with Ponceau staining as a loading control. Band B and Band C indicate immature and mature forms of ΔF508-CFTR, respectively. Band B levels in WT and DKO cells were quantified and expressed as a percentage of siNC levels (right, n=4). Each biological replicate (n) is color-coded. (D) PM expression levels of ΔF508-CFTR-Nluc(Ex) in RNF5/185 DKO cells transfected with 10 nM single siRNA (n=3). (E) Western blot confirmation of AMFR KD in 293MSR WT and RNF5/185 DKO cells, transfected as in panel D. Statistical significance was determined using one-way ANOVA (B, D) with Dunnett’s multiple comparison tests or a two-tailed paired Student’s t-test (C). Data distribution was assumed to be normal but was not formally tested. Data represent mean ± SD. *P < 0.05, **P < 0.01, ***P < 0.0001.

To confirm the on-target effects of AMFR KD, we tested the effect using individual siRNAs. The Nluc assay showed that AMFR KD, using either siAMFR-1 or siAMFR-4, significantly increased the PM level of ΔF508-CFTR in RNF5/185 DKO cells (Fig. 1D). Western blot analysis further confirmed this AMFR KD effect (Fig. 1E).

### AMFR facilitates the ERAD of misfolded CFTR in RNF5/185 DKO cells

To investigate whether AMFR facilitates the ERAD of ΔF508-CFTR, particularly in the absence of RNF5/185, we performed HiBiT-based ERAD kinetic measurements.^29,34^ In contrast to a previous study,^31^ AMFR KD marginally reduced ΔF508-CFTR ERAD in WT cells (Fig. 2A). As previously reported,^29^ ERAD of ΔF508-CFTR was markedly reduced in RNF5/185 DKO cells compared to WT cells (Fig. 2A). Notably, while AMFR KD had little effect in WT cells, it significantly inhibited ERAD in RNF5/185 DKO cells (Fig. 2A). We confirmed the KD of endogenous AMFR protein in the siRNA-transfected cells (Fig. 2B). Similar results were observed in a different cell culture model, BEAS-2B cells derived from human airway epithelial cells. In BEAS-2B cells, AMFR KD significantly reduced the ERAD rate of ΔF508-CFTR-Nluc(CT) only when both RNF5 and RNF185 were knocked down (Fig. 2C and 2D). Consistent with the findings from the HiBiT and Nluc ERAD assays, Western blotting combined with traditional CHX chase showed that AMFR KD dramatically inhibited the elimination of immature ΔF508-CFTR only in the RNF5/185 DKO cells (Fig. 2E). Given recent studies indicating that AMFR participates not only in ERAD but also in the ER-phagy pathway,^35,36^ we investigated whether AMFR-dependent degradation in RNF5/185 DKO cells is affected by proteasome inhibition (bortezomib, BTZ) or lysosome inhibition (bafilomycin A1, BafA1). Since BTZ dramatically inhibited ΔF508-CFTR ERAD and BafA1 treatment partially inhibited ΔF508-CFTR ERAD, it is likely that CFTR is primarily degraded by the proteasome and partially by the lysosome, even in RNF5/185 DKO cells (Fig. 2F). BTZ treatment offset the inhibitory effect of AMFR KD on ΔF508-CFTR ERAD, whereas BafA1 treatment did not show any interaction with AMFR KD effects (Fig. 2F). These findings suggest that AMFR is involved in proteasome-mediated ERAD but not in lysosome-mediated ER-phagy in RNF5/185 DKO cells.

**Figure 2.**
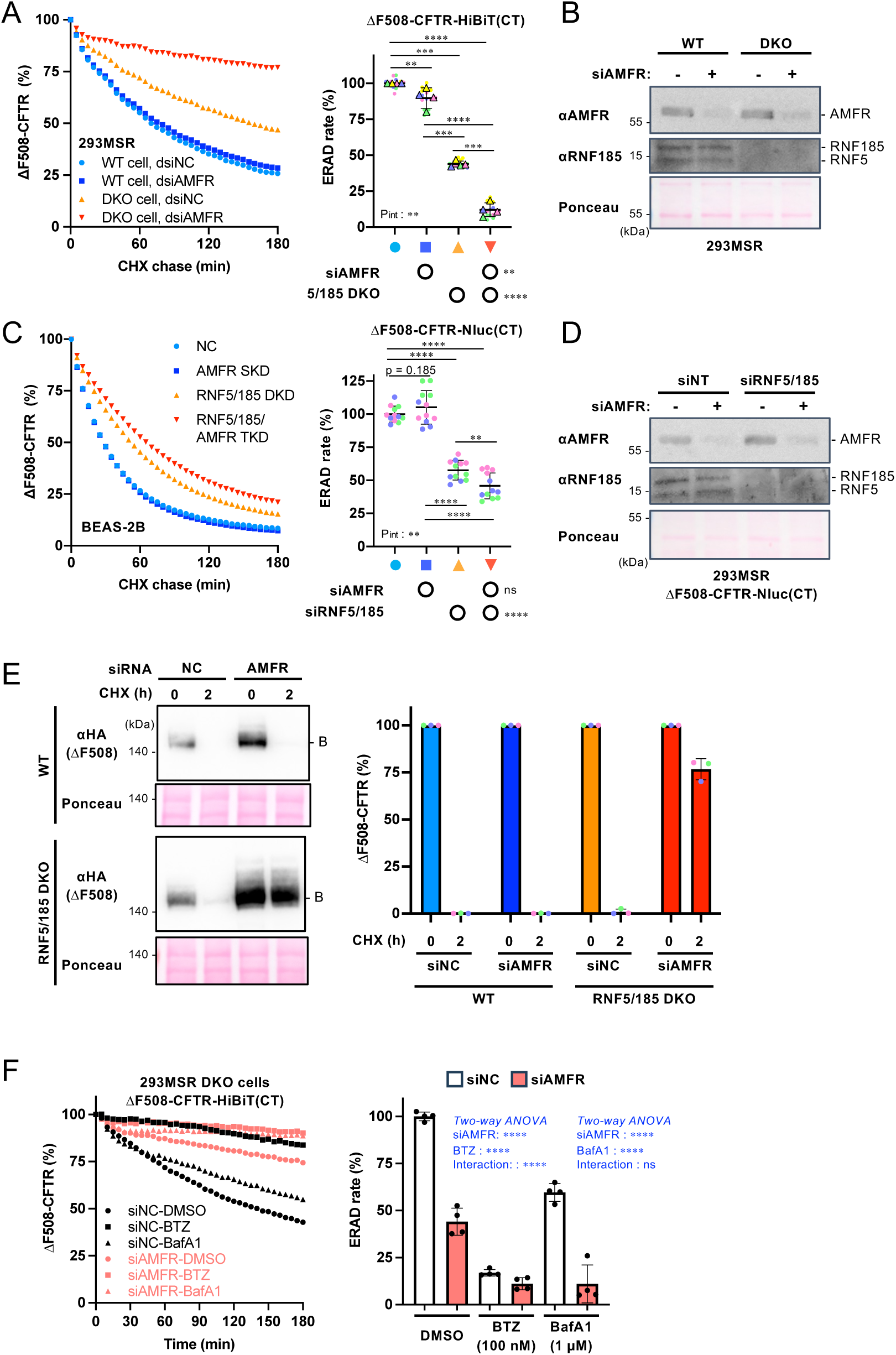
AMFR facilitates the ERAD of ΔF508-CFTR in RNF5/185 DKO cells. (A) Kinetic degradation analysis of ΔF508-CFTR-HiBiT(CT) in 293MSR WT and RNF5/185 DKO cells transfected with 10 nM siNC or pooled siAMFR. Luminescence was continuously measured over 180 min in the presence of CHX and normalized to untreated cells. The ERAD rate of ΔF508-CFTR-HiBiT(CT) was calculated by fitting the initial degradation phase of each kinetic curve (right, n=4). Each biological replicate (n) is color-coded, with averages from four technical replicates represented as triangles. Statistical significance was assessed using two-way RM ANOVA with Holm–Sidak multiple comparison tests, which revealed a significant main effect of AMFR KD or RNF5/185 DKO, as well as a significant interaction between them (P_int_ < 0.05). (B) Western blot confirmation of AMFR KD in 293MSR WT or RNF5/185 DKO cells transfected as in panel A. (C) Kinetic degradation analysis of ΔF508-CFTR-Nluc(CT) in BEAS-2B cells (n=12) transfected with 10 nM pooled siAMFR and/or siRNF5/185 (50 nM pooled siRNF5 + 50 nM pooled siRNF185). The ERAD rate of ΔF508-CFTR-Nluc(CT) was calculated as in panel A. Each independent experiment, consisting of four biological replicates (n), is color-coded. Two-way ANOVA with Holm–Sidak multiple comparison tests revealed a significant main effect of RNF5/185 KD, but not AMFR KD, and a significant interaction between them (P_int_ < 0.05). (D) Western blot confirmation of AMFR and RNF5/185 KD in BEAS-2B cells transfected as in panel C. (E) Western blot analysis of CHX chase measuring ΔF508-CFTR ERAD in 293MSR WT or RNF5/185 DKO HBH-ΔF508-CFTR-3HA cells transfected with 10 nM siNC or pooled siAMFR. Band B (immature ΔF508-CFTR) levels in WT and DKO cells were quantified and expressed as % of T-0 (right, n=3). Each biological replicate (n) is color-coded. (F) ΔF508-CFTR-HiBiT(CT) ERAD in RNF5/185 DKO cells transfected with 10 nM siNC or siAMFR-1, measured in the presence of 0.1% DMSO, 100 nM bortezomib (BTZ), or 1 µM bafilomycin A1 (BafA1). The ERAD rate was calculated by curve fitting of each kinetic degradation curve (right, n=4). Two-way ANOVA revealed a significant main effect of AMFR KD and BTZ or BafA1, and a significant interaction between AMFR KD and BTZ, but not AMFR KD and BafA1 treatment. Data distribution was assumed to be normal but was not formally tested. Data represent mean ± SD. *P < 0.01, ***P < 0.001, ***P < 0.0001, ns (not significant).

### AMFR facilitates the retrotranslocation of misfolded CFTR in RNF5/185 DKO cells

To investigate how AMFR facilitates CFTR ERAD, we examined its impact on retrotranslocation using cytosolic LgBiT and ΔF508-CFTR-HiBiT(Ex), where the HiBiT tag was inserted into the extracellular region of CFTR.^29^ The HiBiT retrotranslocation assay showed that AMFR KD modestly reduced CFTR retrotranslocation in 293MSR WT cells (Fig. 3A). Consistent with our previous findings,^29^ RNF5/185 DKO cells exhibited a robust reduction in the CFTR retrotranslocation, which was further slowed by AMFR KD (Fig. 3A). Similar results were observed using the HiBiT ER disappearance assay, which involved ER-localized LgBiT and ΔF508-CFTR-HiBiT(Ex).^29^ Regardless of RNF5/185 presence, AMFR KD reduced the disappearance of ΔF508-CFTR-HiBiT(Ex) from the ER lumen (Fig. 3B).

**Figure 3.**
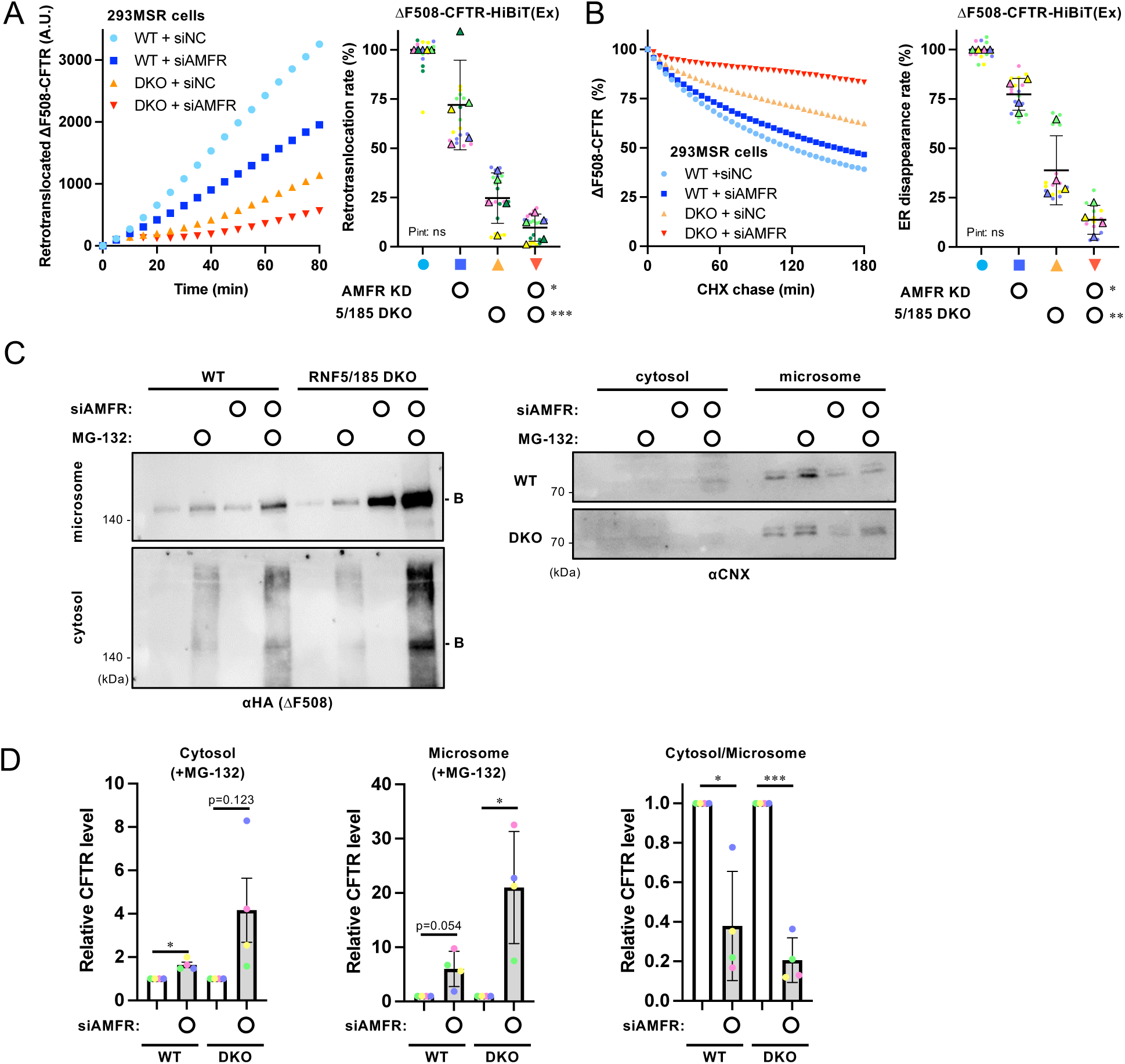
AMFR facilitates the retrotranslocation of ΔF508-CFTR in RNF5/185 DKO cells. (A) Kinetic retrotranslocation analysis of ΔF508-CFTR-HiBiT(Ex) in 293MSR WT and RNF5/185 DKO cells transfected with 10 nM siNC or pooled siAMFR. Luminescence was continuously monitored over 80 min in the presence of 10 µM MG-132, and the signal increase upon MG-132 treatment was plotted as retrotranslocated CFTR. The retrotranslocation rate of ΔF508-CFTR-HiBiT(Ex) was calculated by linear fitting (right, n=5). Two-way RM ANOVA revealed a significant main effect of AMFR KD and RNF5/185 DKO, but no significant interaction between them (P_int_ > 0.05). (B) Kinetic ER disappearance analysis of ΔF508-CFTR-HiBiT(Ex) in 293MSR WT and RNF5/185 DKO cells transfected with 10 nM siNC or pooled siAMFR. Luminescence was continuously monitored over 180 min in the presence of CHX and normalized to untreated cells to represent remaining CFTR at the ER (%). The ER disappearance rate of ΔF508-CFTR-HiBiT(Ex) was calculated by fitting the kinetic ER disappearance curve (right, n=4). Two-way RM ANOVA with Holm–Sidak multiple comparison tests revealed a significant main effect of AMFR KD and RNF5/185 DKO, but no significant interaction between them (P_int_ > 0.05). (C, D) Western blot analysis of CFTR levels in microsomal and cytosolic fractions of 293MSR WT and RNF5/185 DKO HBH-ΔF508-CFTR cells transfected with 10 nM siNC or pooled siAMFR. Cells were treated with or without 10 µM MG-132 for 3 hours before subcellular fractionation. Microsome enrichment (ER membranes) was confirmed using an anti-calnexin (CNX) antibody (C, right). The quantities of cytosolic and ER-localized ΔF508-CFTR were determined by densitometry, and the ratio of cytosolic to ER-resident CFTR was quantified as a measure of retrotranslocation efficiency (D, n=4). Statistical significance was assessed using a two-tailed paired t-test. Each biological replicate (n) is color-coded. Data distribution was assumed to be normal but was not formally tested. Data represent mean ± SD. *P < 0.05, **P < 0.01, ***P < 0.001, ns (not significant).

To confirm these findings, we performed subcellular fractionation, using calnexin (CNX) as a marker for the ER membrane. Western blotting showed that cytosolic ΔF508-CFTR appeared as smear bands only after proteasome inhibition with MG-132 (Fig. 3C). These smear bands likely represent ubiquitinated CFTR that underwent retrotranslocation and subsequent proteasomal degradation.^37,38^ AMFR KD increased ΔF508-CFTR in the cytosol, but also resulted in an increase in CFTR levels in the microsomes in WT and RNF5/185 DKO cells (Fig. 3D). Quantitative analysis of the cytosol-to-microsome ratio of ΔF508-CFTR revealed that AMFR KD predominantly reduced the cytosolic fraction relative to microsomes, supporting the notion that AMFR facilitates CFTR retrotranslocation even in the absence of RNF5/185 (Fig. 3D).

### AMFR reduces the ubiquitination of misfolded CFTR in RNF5/185 DKO cells

AMFR has been proposed to function as an E4-like enzyme, facilitating CFTR ubiquitination by extending polyubiquitin (polyUb) chains initiated by the E3 ligase RNF5.^31^ To determine whether AMFR regulates CFTR ubiquitination even in the absence of RNF5/185, we assessed CFTR ubiquitination using multiple methodologies. Western blotting of isolated HBH-ΔF508-CFTR under denaturing conditions by Neutravidin (NA) pull-down showed that AMFR KD reduced pan-ubiquitination in both WT and RNF5/185 DKO cells (Fig. 4A and 4B). Additionally, Ub ELISA^39^ using a K48-linked polyUb-specific antibody revealed that AMFR KD reduced CFTR K48-linked polyubiquitination in WT cells (Fig. 4C). While its effect was attenuated in RNF5/185 DKO cells, AMFR KD still showed a tendency to reduce K48-linked polyubiquitination (Fig. 4C). To further investigate the role of AMFR in polyUb chain formation, we analyzed its impact on various polyUb chain configurations using ELISA with Myc-Ub mutants, each containing a single lysine (K) residue. The results showed that AMFR KD tended to decrease all tested polyUb chain types, including K29-, K33-, and K63-linked polyUb chains on CFTR in RNF5/185 DKO cells (Fig. 4D). These findings suggest that AMFR promotes diverse forms of CFTR polyubiquitination, even in the absence of RNF5/185, highlighting its role in ERAD regulation.

**Figure 4.**
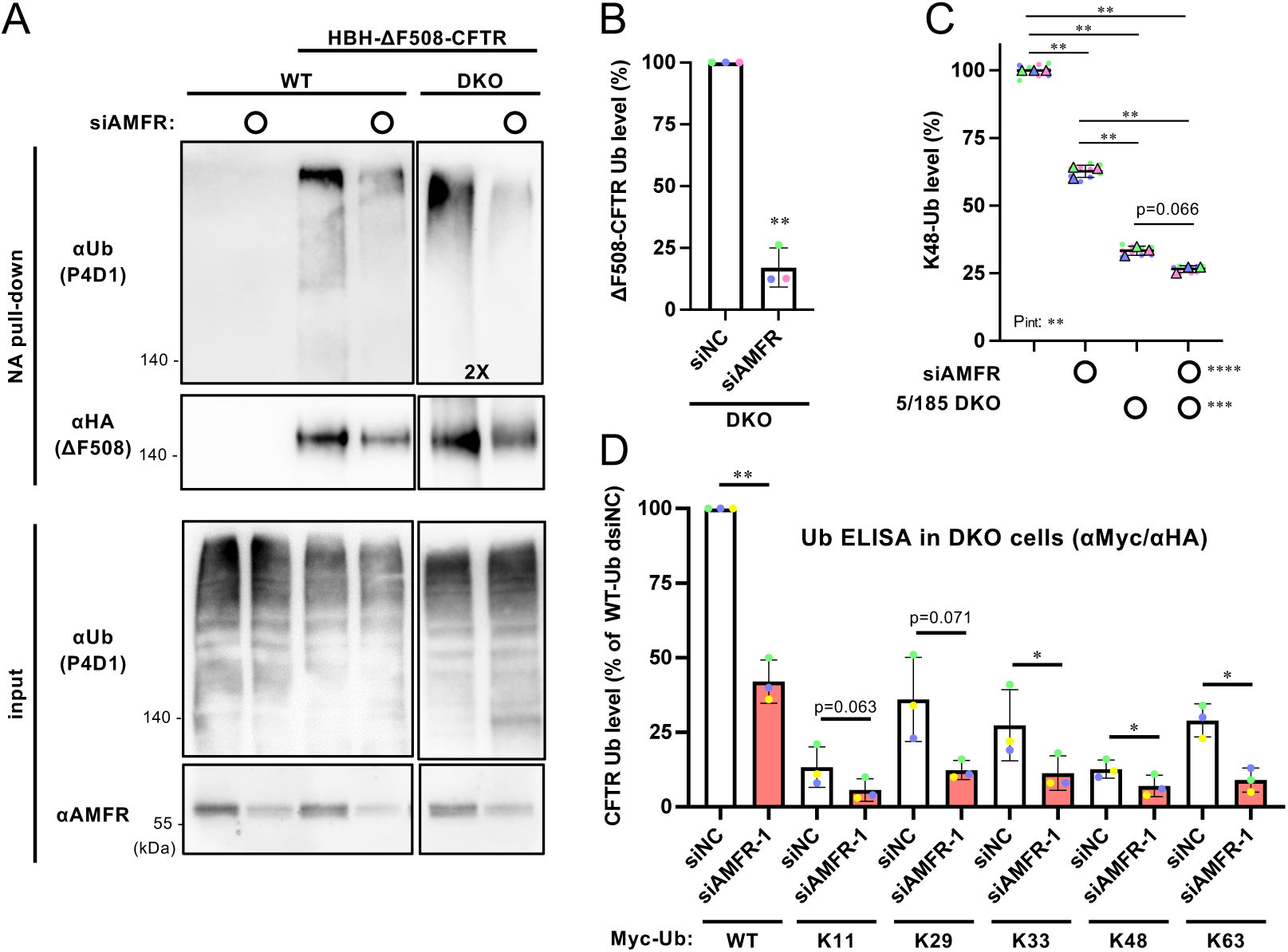
AMFR facilitates the ubiquitination of ΔF508-CFTR in RNF5/185 DKO cells. (A, B) Ubiquitination levels of ΔF508-CFTR in HBH-ΔF508-CFTR-3HA 293MSR WT and RNF5/185 DKO cells transfected with 10 nM siNC or pooled siAMFR were measured by NA pull-down under denaturing conditions, followed by Western blot analysis. To enhance detection of CFTR ubiquitination in DKO cells, a double amount of precipitate sample (2×) compared to WT cells was used. The CFTR ubiquitination level in RNF5/185 DKO cells was quantified by densitometry and normalized to CFTR levels in the precipitates (B). Statistical significance was assessed using a two-tailed paired t-test (n=3). (C) K48-linked polyubiquitination of ΔF508-CFTR in HBH-ΔF508-CFTR-3HA 293MSR WT and RNF5/185 DKO cells transfected with 10 nM siNT or pooled siAMFR was quantified by Ub ELISA using a K48-linkage-specific polyUb antibody. Cells were treated with 10 µM MG-132 for 3 hours at 37°C. The ubiquitination level was normalized to the CFTR amount quantified by ELISA using an anti-HA antibody (n=3). Two-way RM ANOVA with Holm–Sidak multiple comparison tests revealed significant main effects of AMFR KD and RNF5/185 DKO, as well as a significant interaction between them (P_int_ < 0.01). (D) Analysis of various polyUb linkages on ΔF508-CFTR in RNF5/185 DKO HBH-ΔF508-CFTR-3HA cells transfected with Myc-Ub variants and 10 nM siNC or siAMFR-1. Ub ELISA was performed using anti-Myc and anti-HA antibodies (n=3). The Ub levels were expressed as a percentage relative to the siNC and Myc-WT Ub-transfected sample. Each biological replicate (n) is color-coded. Statistical significance was assessed using a two-tailed paired t-test. Data distribution was assumed to be normal but was not formally tested. Data represent mean ± SD. *P < 0.05, **P < 0.01.

### SYVN1 is not involved in the CFTR ERAD in RNF5/185 DKO cells

SYVN1 is a mammalian homolog of yeast Hrd1 and shares amino acid sequence homology with AMFR.^40,41^ However, despite their similarity, SYVN1 and AMFR perform distinct cellular functions.^31,42^ To determine whether SYVN1 also plays a role in the ERAD bypass pathway in RNF5/185 DKO cells, we examined the impact of SYVN1 KD on ΔF508-CFTR ERAD. Unlike AMFR, SYVN1 KD failed to inhibit ΔF508-CFTR-HiBiT(CT) degradation in both WT and RNF5/185 DKO cells (Fig. 5A). The efficiency of SYVN1 KD using two different siRNAs was confirmed by Western blotting (Fig. 5B). Consistent with the ERAD assay results, SYVN1 KD had a weaker effect on ΔF508-CFTR protein levels compared to AMFR KD (Fig. 5B). Furthermore, unlike AMFR, SYVN1 KD failed to increase the cell surface expression of ΔF508-CFTR-Nluc(Ex) in RNF5/185 DKO cells (Fig. 5C). However, pulldown experiments showed that both AMFR and SYVN1 physically interact with immature ΔF508-CFTR in the ER of WT and RNF5/185 DKO cells, despite their differing effects on CFTR ERAD (Fig. 5D). Ub ELISA revealed that SYVN1 KD reduced K48-linked polyubiquitination of ΔF508-CFTR in 293MSR WT cells, despite having no impact on ERAD (Fig. 5E). However, in RNF5/185 DKO cells, SYVN1 KD failed to reduce CFTR K48-linked polyubiquitination (Fig. 5E).

**Figure 5.**
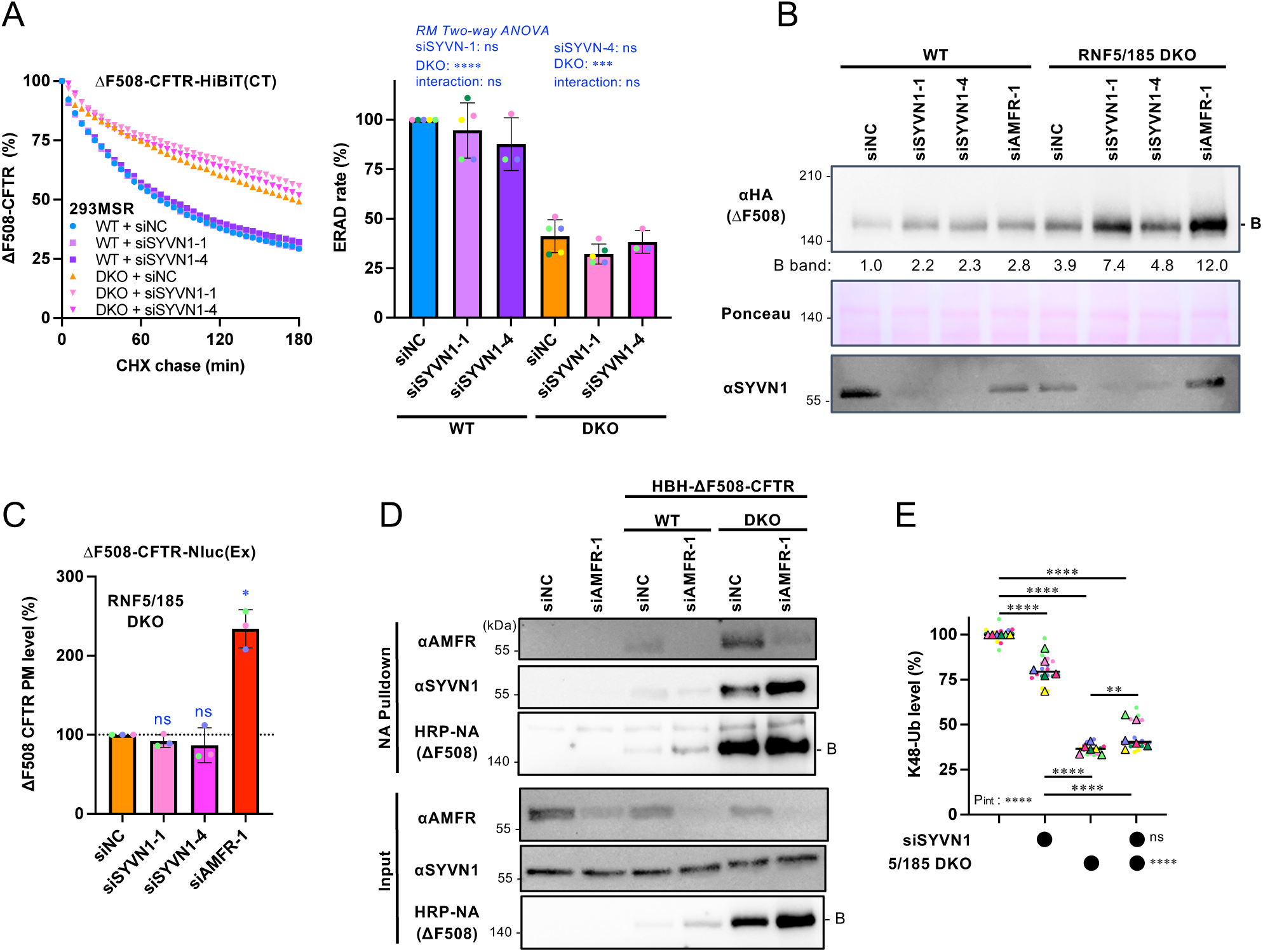
The minimal role of SYVN1 in CFTR ERAD bypass in RNF5/185 DKO cells. (A) Kinetic degradation of ΔF508-CFTR-HiBiT(CT) in 293MSR WT and RNF5/185 DKO cells transfected with 10 nM siNT, siSYVN1-1, or siSYVN1-4 was assessed as described in Figure 2A. The ERAD rate was calculated by fitting the initial degradation portion of each kinetic degradation curve (right, n=3–5). Each biological replicate (n) is color-coded, with the averages from three to four technical replicates represented by triangles. Statistical significance was determined using two-way RM ANOVA with Holm–Sidak multiple comparison tests. (B) Western blot analysis of HBH-ΔF508-CFTR-3HA levels in 293MSR WT and RNF5/185 DKO cells transfected as in panel A. Ponceau staining was used as a loading control. The relative levels of immature ΔF508-CFTR (band B) were quantified via densitometry and are indicated below the anti-HA blot. SYVN1 KD was confirmed using an anti-SYVN1 antibody. (C) PM expression levels of ΔF508-CFTR-Nluc(Ex) in RNF5/185 DKO cells transfected with 10 nM of the indicated siRNA (n=3). Statistical significance was evaluated using one-way RM ANOVA with Dunnett’s multiple comparison tests. (D) Interaction of HBH-ΔF508-CFTR with endogenous AMFR or SYVN1 in 293MSR WT or RNF5/185 DKO cells transfected with 10 nM siNC or siAMFR-1, analyzed by NA pull-down and Western blotting. (E) K48-linked polyubiquitination levels of ΔF508-CFTR in HBH-ΔF508-CFTR-3HA 293MSR WT or RNF5/185 DKO cells transfected with 10 nM siNC or pooled siSYVN1 (siSYVN1-1 and siSYVN1-4) were quantified via Ub ELISA using an anti-K48-Ub antibody. Cells were treated with 10 µM MG-132 for 3 hours before lysis. Ub levels were normalized to CFTR expression, quantified via ELISA using an anti-HA antibody (n=6). Statistical significance was assessed using two-way RM ANOVA with Holm–Sidak multiple comparison tests. Each biological replicate (n) is color-coded. Statistical significance was determined using a two-tailed paired t-test. Data are presented as mean ± SD. *P < 0.05, **P < 0.01, ***P < 0.001, ****P < 0.0001, ns (not significant).

### The AMFR-mediated ERAD bypass is involved in the ERAD of Insig-1 and TCRα

To better understand the general role of AMFR-mediated ERAD bypass in ERQC system, we examined the impact of AMFR KD in RNF5/185 DKO cells on the ERAD of various membrane proteins (Fig. 6A). ΔY490-ABCB1 (MDR1/P-glycoprotein) is an ERAD-C substrate analogous to ΔF508-CFTR^43^ and is degraded via the RNF5/185 ERAD pathway. ^29^ TCRα and Insig-1 have been classified as ERAD-Lm^44^ and ERAD-M substrates,^18,45^ respectively. The HiBiT degradation assay revealed that AMFR KD significantly inhibited the ERAD of Insig-1, but had only a marginal effect on TCRα ERAD in 293MSR WT cells (Fig. 6B, 6C). Consistent with previous studies,^29^ ERAD of Insig-1 and TCRα was unexpectedly enhanced in RNF5/185 DKO cells, possibly due to compensatory mechanisms (Fig. 6B, 6C). Notably, the effect of AMFR KD on ERAD of Insig-1 and TCRα was significantly stronger in RNF5/185 DKO cells, leading to robust inhibition of their degradation (Fig. 6B, 6C). Surprisingly, AMFR KD also significantly reduced the ERAD of ΔY490-ABCB1 in WT cells, but its impact was diminished in RNF5/185 DKO cells (Fig. 6D). The AMFR-mediated ERAD branch also contributed to the degradation of N1303K-CFTR, a misfolded CFTR mutant with an NBD2 mutation that disrupts the membrane-spanning domains (MSD1 and MSD2).^46^ N1303K-CFTR has been reported to be degraded through both ERAD and ER-phagy pathways.^47^ AMFR KD significantly reduced N1303K-CFTR ERAD in WT cells, but its effect was less pronounced in RNF5/185 DKO cells (Fig. 6E).

**Figure 6.**
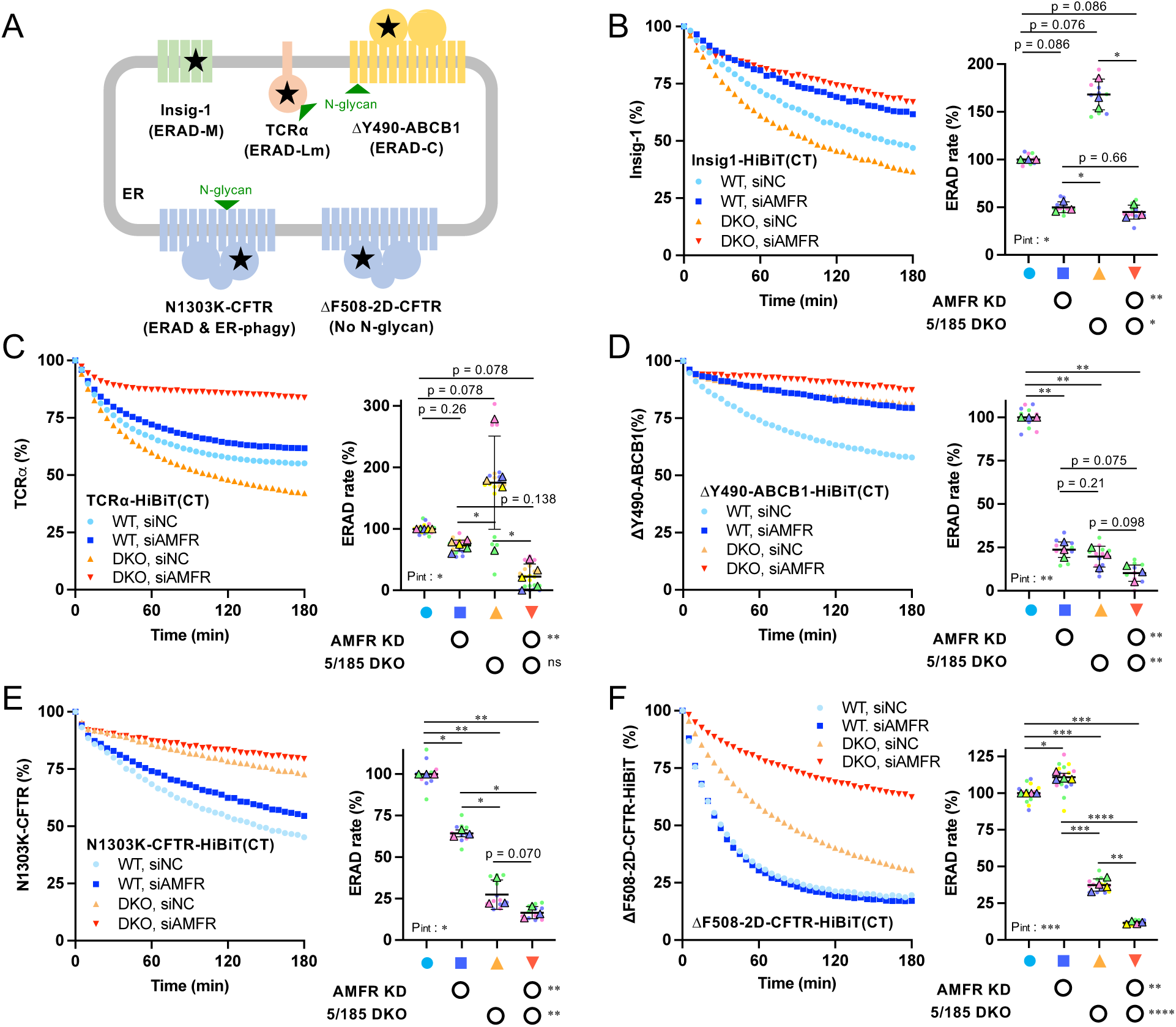
Contribution of the AMFR-mediated ERAD bypass to other ERAD substrates. (A) Schematic representation of the ERAD substrate models. The misfolded regions are marked with stars, and the HiBiT tag was fused to the cytoplasmic domain. N-glycans are indicated for glycoproteins. (B-F) The HiBiT degradation assay was used to assess the ERAD of Insig1-HiBiT (B, n=3), TCRα-HiBiT (C, n=5), ΔY490-ABCB1-HiBiT (D, n=3), N1303K-CFTR-HiBiT (E, n=3), and ΔF508-2D-CFTR-HiBiT (F, n=4) in 293MSR WT and RNF5/185 DKO cells transfected with 10 nM siNC or siAMFR-1, as described in Figure 2A. Each biological replicate (n) is color-coded, with the averages from four technical replicates represented by triangles. Statistical significance was determined using two-way RM ANOVA with Holm–Sidak multiple comparison tests. The analysis identified a significant main effect of AMFR KD or RNF5/185 DKO (except for panel C) and an interaction between them (Pint < 0.05). Data are presented as mean ± SD. *P < 0.05, **P < 0.01, ***P < 0.001, ns (not significant).

To determine whether the AMFR-mediated ERAD branch is involved in the N-glycan-dependent ERAD (GERAD) pathway,^48,49^ we tested ΔF508-2D-CFTR, in which two asparagine residues (N894, N900) were mutated to aspartic acid (N894D, N900D) to abolish N-glycosylation.^50^ However, GERAD does not appear to be involved in the AMFR-mediated ERAD pathway, as AMFR KD had an equivalent effect on the ERAD of ΔF508-2D-CFTR and ΔF508-CFTR in both WT and RNF5/185 DKO cells (Fig. 6F). To further investigate which regions of misfolded CFTR are recognized by the AMFR-mediated ERAD pathway, we analyzed isolated CFTR domains. These isolated domains possess intrinsic conformational defects^46^ and are typically targeted for degradation by cellular protein QC mechanisms.^29^ The HiBiT assay demonstrated that AMFR KD had minimal impact on the ERAD of CFTR MSD1 and MSD2 in WT cells. In contrast, in RNF5/185 DKO cells, AMFR KD significantly impaired the degradation of these domains (Fig. 7A and 7B). However, AMFR KD did not significantly affect the degradation of the ΔF508-CFTR NBD1 (Fig. 7C). To further assess the physical interaction between AMFR and CFTR domains, we performed pull-down experiments in RNF5/185 DKO cells expressing HiBiT-tagged CFTR domains and AMFR fused to a myc-biotin (MB) tag.^51^ Robust binding of AMFR was observed with CFTR MSM1 and MSD2, whereas its interaction with NBD1 was relatively weak (Fig. 7D). Interestingly, high molecular weight smear bands were also detected in the AMFR precipitates (Fig. 7D), suggesting that these bands may represent ubiquitinated CFTR forms. These findings suggest that AMFR preferentially recognizes CFTR transmembrane domains and facilitates ERAD bypass in the absence of RNF5/185.

**Figure 7.**
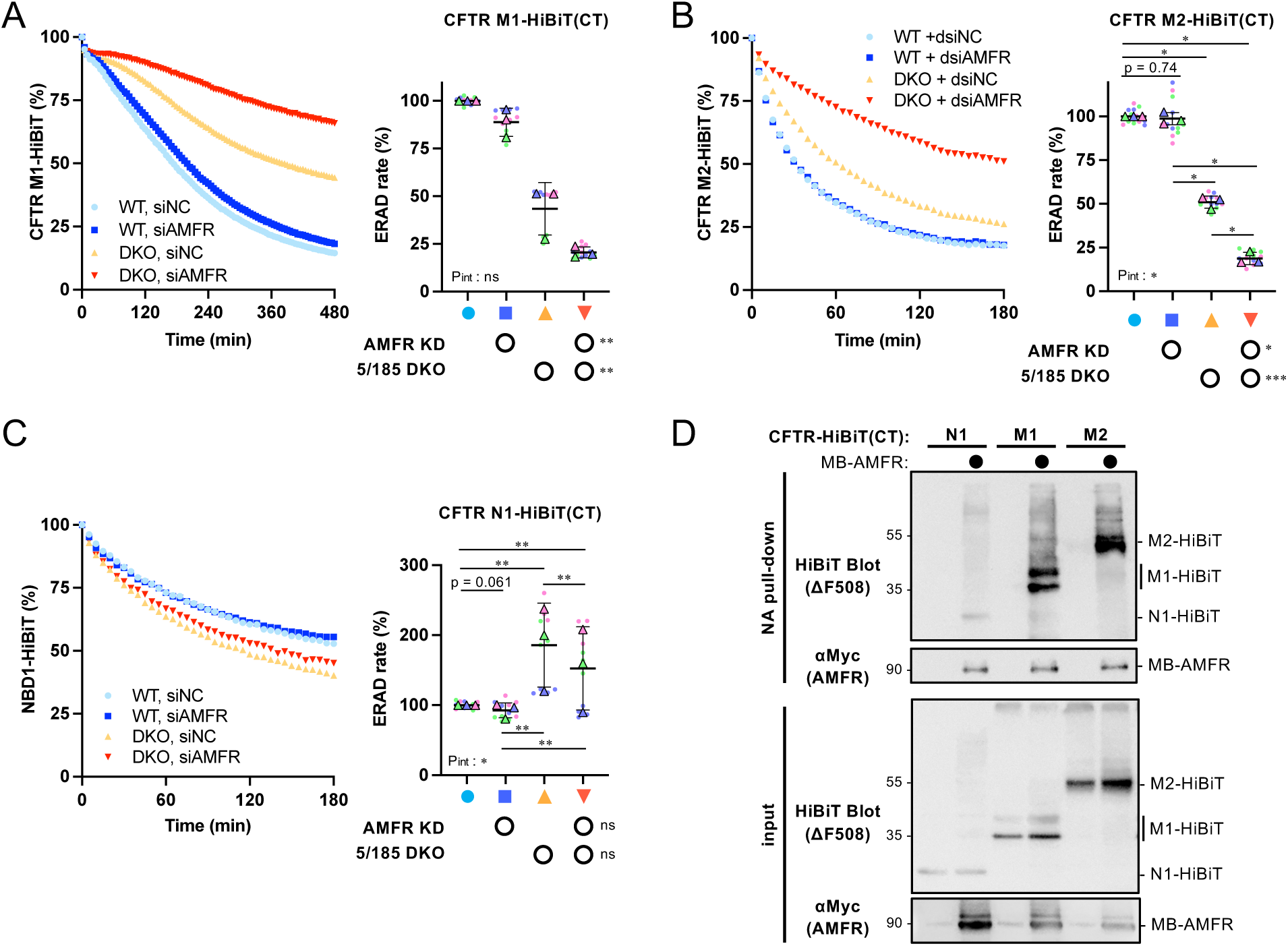
AMFR recognizes and facilitates ERAD of CFTR transmembrane domains in RNF5/185 DKO cells. (A-C) The HiBiT degradation assay was used to assess the ERAD of CFTR MSD1 (M1)- HiBiT (A, n=3), MSD2 (M2)-HiBiT (B, n=3), and NBD1 (N1)-HiBiT (C, n=3) in 293MSR WT and RNF5/185 DKO cells transfected with 10 nM siNC or siAMFR-1, as described in Figure 2A. Each biological replicate (n) is color-coded, with the averages from four technical replicates represented by triangles. Statistical significance was determined using two-way RM ANOVA with Holm–Sidak multiple comparison tests. Data are presented as mean ± SD. *P < 0.05, **P < 0.01, ***P < 0.001, ns (not significant). (D) The interaction between MB-AMFR and co-expressed CFTR N1-, M1-, and M2-HiBiT in RNF5/185 DKO cells was analyzed by NA pull-down followed by Western blotting. Cells were pre-treated with 10 µM MG-132 for 3 hours at 37°C before lysis. The HiBiT-fused CFTR domain proteins were detected using the HiBiT Blotting system, while MB-AMFR was detected with an anti-Myc antibody.

## Discussion

This study demonstrates that AMFR compensates for CFTR ERAD when the major RNF5/185-dependent ERAD pathway is impaired. Notably, this complementary effect was not observed with SYVN1, which is involved in the ERAD-L and ERAD-M pathways.^6,15–17^ This suggests that the effect of AMFR is not a general property of ER-localized E3 ligases but rather a specific compensatory mechanism. Different functions of AMFR and SYVN1 have been reported, and our findings are in agreement with these previous studies.^52,53^ Although AMFR is primarily involved in the ERAD-M pathway, it may compensate for the dysfunction of the ERAD-C pathway, typically managed by RNF5/185, when required. In fact, AMFR expression is upregulated under ER stress, a condition in which aberrant ER proteins accumulate.^54^ In RNF5/185-deficient cells, ER stress could be induced, leading to increased AMFR expression, which may enable this compensatory role. Indeed, ER stress markers including DDIT3 (CHOP) and ERN1 (IRE1) are highly expressed in RNF5/185 DKO cells (Fig. S2). Since AMFR is known to mitigate ER stress,^55^ the ERAD bypass pathway may aid in the degradation of diverse misfolded proteins, thereby maintaining ER proteostasis. Supporting this model, a recent study suggests that AMFR may play an increased role in ΔF508-CFTR degradation in RNF5 KO cells.^32^ Additionally, the degradation of TAP2 in RNF5/185 DKO cells was found to be inhibited by AMFR KD.^56^ These findings suggest that AMFR plays a crucial role in the ERAD bypass pathway, particularly when the RNF5/185-mediated ERAD branch is compromised. This highlights its involvement in the degradation of a variety of misfolded proteins, extending beyond CFTR.

Although ΔF508-CFTR is generally classified as an ERAD-C substrate,^25,26^ its potential structural abnormalities in the transmembrane domains cannot be ruled out, given that ΔF508-CFTR misfolding correctors are known to bind to the transmembrane domains.^57^ Indeed, structural defects in the CFTR MSDs have been identified through in situ CFTR trypsin sensitivity assays.^46^ Furthermore, our findings that AMFR regulates the ERAD of isolated CFTR MSD1/2 further support the idea that ΔF508-CFTR may exhibit structural abnormalities in both cytosolic and transmembrane regions. Indeed, it has been suggested that RNF185 may recognize the transmembrane domain of CYP51A1 and facilitate its ERAD.^58^ The CFTR-RNF5-RNF185 complex predictions from AlphaFold3 suggest that RNF5 and RNF185 may interact with the transmembrane region of ΔF508-CFTR, especially transmembrane segment 1 (TM1) (Fig. S3A), providing support for the notion that RNF5/185 is involved in the ERAD-M pathway rather than the ERAD-C pathway. Thus, RNF5/185 may regulate CFTR degradation through both the ERAD-M and ERAD-C pathways, resembling the dual role of Doa10, a yeast E3 ligase that participates in both ERAD-M and ERAD-C branches.^6,11,16,17^ A prediction from AlphaFold3 of the CFTR-AMFR complex suggests that AMFR also recognizes the CFTR TM1, with its binding site partially overlapping with those of RNF5 and RNF185 (Fig. S3B, S3C). Thus, AMFR may be capable of recognizing the transmembrane domain of CFTR, potentially compensating for the function of RNF5/185 when its activity is compromised or insufficient.

Although AMFR has previously been reported to contribute to CFTR ERAD,^31^ our highly quantitative HiBiT-based ERAD assay indicates that its role under normal conditions is limited across multiple cell culture models. A prior study proposed that AMFR functions as an E4-like enzyme downstream of RNF5.^31^ However, our results demonstrate that AMFR has an RNF5/185-independent function in the ERAD of membrane proteins. Since AMFR KD reduces CFTR ubiquitination levels in WT cells, the possibility that AMFR functions as an E4-like enzyme in the presence of RNF5/185 cannot be excluded. While AMFR-mediated ubiquitination appears dispensable for efficient CFTR ERAD in WT cells, it facilitates diverse polyubiquitin modifications— such as K29- and K63-linked chains—in RNF5/185 DKO cells. These modifications likely contribute to CFTR degradation through an ERAD bypass pathway, in which alternative ubiquitin linkages may play a compensatory role. Although AMFR has also been implicated in ER-phagy,^35,36^ our results indicate that this ERAD bypass pathway is insensitive to lysosome inhibitors but is suppressed by proteasome inhibitors, confirming that it functions as part of the ERAD pathway rather than ER-phagy.

The AMFR-dependent ERAD bypass pathway plays a lesser role in the degradation of relatively stable RNF5/185-dependent ERAD substrates, such as ΔY490-ABCB1 and N1303K-CFTR, compared to ΔF508-CFTR. This suggests that the multilayered ERAD bypass mechanism may not be involved in the degradation of substrates with mild conformational abnormalities and relatively slow degradation rates. However, more severely misfolded proteins, which could lead to cytotoxicity, may require a multilayered ERAD branch for efficient degradation, helping maintain proteostasis. While the physiological function of the AMFR ERAD bypass remains unclear, it may become particularly relevant in pathological environments, such as myocardial infarction and ischemia,^59^ where RNF5 expression is reduced.

Unlike in yeast, the ERAD branches in mammalian cells are highly diverse and challenging to analyze due to functional overlap.^2,8^ Complementary functions are likely to become more pronounced in cases of long-term functional decline, such as in the ERAD-associated Ub ligase KO cells, which can aid in the analysis of more redundant and complementary ERAD branch mechanisms. The use of multiple Ub ligases KO and/or KD, combined with highly quantitative ERAD quantification in this study is expected to deepen our understanding of the molecular mechanisms underlying the independence and complementation of ERAD branches in mammalian cells.

## Acknowledgments

We thank R. Kopito and Y. Ye for the TCRα-HA, N. Hosokawa for the mouse AMFR, and D. Root (Addgene #41394 pLIX402, #25890 pLX304) for providing expression vectors. We appreciate the RNA-seq analysis support from Y. Mizukami and K. Watanabe (Yamaguchi University), and we thank M. Okumura and S. Kanemura (Tohoku University) for valuable discussions regarding the structural data.

## Author Contributions

Conceptualization, T.O.; methodology, U.N., A.F., Y.K.; investigation, U.N., A.F., S.K.; data curation, U.N., A.F.; formal analysis, T.O.: writing – original draft, U.N., A.F., T.O.; writing – review & editing, Y.K., T.O.; visualization, T.O.: funding acquisition, Y.K., T.O.; resources, T.O.; supervision, T.O.

## Lead contact

Further information and requests for resources and reagents may be directed to and will be fulfilled by Tsukasa Okiyoneda.

## Data Availability

The raw data required to reproduce these findings are available from the corresponding authors upon reasonable request.

### Declaration of generative AI and AI-assisted technologies in the writing process

During the preparation of this work the authors used ChatGPT in order to improve language and readability. After using this tool/service, the authors reviewed and edited the content as needed and take full responsibility for the content of the publication.

## Materials and methods

### Plasmids and antibodies

Plasmids, chemicals, and antibodies used in this study are listed in the Key Resources Table. pcDNA3.1(-) Myc-Ubiquitin WT and its single-lysine (K) mutants, in which only one lysine residue remains at the specified position, were constructed using PCR-based cloning methods. pNUT ΔF508-2D-CFTR-3HA-HiBiT(CT) was generated by introducing N-glycosylation site mutations (N894D and N900D) into pNUT ΔF508-CFTR-3HA-HiBiT(CT),^29^ as previously described.^50^ TCRα-HiBiT was constructed by inserting TCRα from pCMV-TCRα-HA (a kind gift from Dr. R. Kopito) into the pBiT2.1-HiBiT vector via In-Fusion cloning. MB-AMFR was created by inserting mouse AMFR from pcDNA3.1 (a kind gift from Dr. N. Hosokawa) into the pBiT2.1-HiBiT vector using Gateway cloning, with pcDNA-Myc-Bio as the Gateway destination vector.^51^

### Cell lines and cell culture

293MSR, RNF5/185 DKO cells, and their respective transfectants were cultured as previously described.^29^ 293MSR and RNF5/185 DKO Tet-on cells stably expressing ΔF508-CFTR-Nluc(Ex) were generated via lentiviral transduction, as previously reported, ^33,39^ and maintained in DMEM medium supplemented with 10% fetal bovine serum (FBS), 0.5 mg/ml G418, and 5 µg/ml blasticidin S. BEAS-2B Tet-on ΔF508-CFTR-3HA-Nluc(CT) cells were cultured following previously established protocols.^29^ CFTR expression in 293MSR and BEAS-2B cells was induced by treatment with 1 µg/ml doxycycline (Dox) for 2 days (293MSR) or 4 days (BEAS-2B), unless otherwise specified. All cell culture media were supplemented with 100 U/ml penicillin and 100 μg/ml streptomycin (FUJIFILM Wako Pure Chemical Corporation), and cells were maintained at 37°C under 5% CO₂.

### Mammalian cell transfection

Transient transfection in 293MSR and RNF5/185 DKO cells was performed using polyethylenimine (PEI) Max (Polysciences Inc.), and experiments were conducted 2 days post-transfection. siRNA transfections (typically at 10 nM) were carried out using Lipofectamine RNAiMax (Invitrogen) following the manufacturer’s instructions. Unless otherwise specified, siRNA-transfected cells were used for experiments 4 days post-transfection. The siRNAs used in this study are listed in the Key Resources Table. Unless explicitly stated, pooled siRNA was used for KD. The pooled siRNA was prepared by mixing equal amounts of two or three individual siRNAs, as listed in the Key Resources Table. Negative control siRNAs included siNC (Integrated DNA Technologies, IDT, Coralville, IA) and AllStars Negative Control siRNA (siNT) (Qiagen, Venlo, The Netherlands).

### Whole-transcriptome analysis with RNA-seq

Total RNA was extracted from 293MSR cells using the RNeasy Mini Kit (Qiagen Inc., Hilden, Germany). Library preparation was performed as previously described,^60,61^ and the quality and concentration of the libraries were assessed using an Agilent 2200 TapeStation (Agilent Technologies Inc.) with a D1000 ScreenTape assay. Libraries were then pooled at equal molecular concentrations and sequenced on an Illumina NextSeq500 DNA sequencer using a 75-bp paired-end cycle sequencing kit (Illumina Inc., San Diego, CA). The sequencing data were trimmed and mapped to the human reference genome (GRCm38 release-95) using the CLC Genomics Workbench software (ver. 12.0.3; Qiagen Inc., RRID:SCR_011853). Mapped read counts were normalized to transcripts per million (TPM) and subsequently log₂-transformed after adding 1.

### PM density of CFTR HiBiT assay

The PM level of ΔF508-CFTR-Nluc(Ex) was measured as previously.^33^ Briefly, ΔF508-CFTR-Nluc(Ex) 293MSR WT or RNF5/185 DKO cells in 96 well white plate were incubated with Extracellular NanoLuc^®^ Substrate (Promega, CS313501, Custom Assay Services) in 50 µl/well CO_2_ independent medium (ThermoFisher) for 5-10 min, then the luminescence was measured by plate readers Luminoskan (ThermoFisher), Varioskan (ThermoFisher), or Enspire (PerkinElmer). The ΔF508-CFTR-Nluc(Ex) expression was induced by 1 µg/ml Dox treatment for 2 days at 37°C.

### Western blotting

Western blotting was performed as previously described.^29^ Briefly, cells were solubilized in RIPA buffer supplemented with 1 mM PMSF (FUJIFILM Wako Pure Chemical Corporation), 5 µg/mL leupeptin (FUJIFILM Wako), and 5 µg/mL pepstatin A (Peptide Institute, Inc.). Equal amounts of cell lysate proteins were subjected to SDS-PAGE, and the proteins were transferred onto nitrocellulose membranes using the Mini Trans-Blot cell system (Bio-Rad, Hercules, USA). Membranes were then blocked with 5% skim milk and incubated with primary antibodies, followed by HRP-conjugated secondary antibodies. Signals were detected using SuperSignal West Pico PLUS Chemiluminescent Substrate (ThermoFisher), ImmunoStar Zeta (FUJIFILM Wako Pure Chemical Corporation), or ImmunoStar LD (FUJIFILM Wako Pure Chemical Corporation). Western blot images were acquired using either the LAS 4000 mini (Cytiva Life Sciences, Marlborough, MA, USA) or the FUSION Chemiluminescence Imaging System (Vilber Bio Imaging, Quantum, France). In some experiments, the HiBiT tag was detected using the Nano-Glo^®^ HiBiT Blotting System (Promega), following the manufacturer’s instructions. For loading control, total proteins were stained with Ponceau S. Densitometric analysis of Western blots was performed using ImageQuant (LAS 4000 mini software, Cytiva) or Evolution-Capt (Fusion software, Vilber Bio Imaging).

### CFTR Ubiquitination Measurement by Western blotting

CFTR ubiquitination in cells was measured as previously described.^29^ Briefly, 293MSR and RNF5/185 DKO cells stably expressing HBH-ΔF508-CFTR-3HA were lysed in RIPA buffer containing 5 µg/mL leupeptin, 5 µg/mL pepstatin A, 1 mM PMSF, 10 µM MG-132, and 5 mM N-ethylmaleimide (NEM) following 10 µM MG-132 treatment at 37 °C for 3 hours. Immature HBH-ΔF508-CFTR was purified under denaturing conditions using NeutrAvidin agarose (ThermoFisher) and analyzed by Western blotting with anti-Ub (P4D1) and anti-HA (16B12, BioLegend) antibodies. The CFTR ubiquitination level was quantified by densitometry and normalized to the CFTR level in the precipitate.

### Ub ELISA

CFTR ubiquitination levels in 293MSR and RNF5/185 DKO Tet-on HBH-ΔF508-CFTR-3HA cells were measured as previously described.^39,62^ Briefly, cells were lysed in RIPA buffer supplemented with 5 µg/mL leupeptin, 5 µg/mL pepstatin A, 1 mM PMSF, 10 µM MG-132, and 5 mM N-ethylmaleimide (NEM) after treatment with 10 µM MG-132 at 37 ° C for 3 hours. HBH-ΔF508-CFTR-3HA in the cell lysate was immobilized onto NeutrAvidin-coated 96-well white plates and denatured in 8 M urea at room temperature for 5 minutes. The plate was then washed five times with 2 M urea in RIPA buffer, followed by four washes with 0.1% NP-40 in PBS. After blocking with 0.1% BSA, ubiquitination and CFTR levels were detected using anti-K48 Ub (Apu2, Merck Millipore) and anti-HA (16B12, BioLegend) antibodies, respectively. CFTR ubiquitination levels were then normalized to the total CFTR level.

To detect Myc-tagged Ub conjugation on CFTR, 293MSR WT and RNF5/185 DKO Tet-on HBH-ΔF508-CFTR-3HA cells were transfected with Myc-Ub WT or Ub variants containing only a single lysine (K) residue at the specified position (K11, K29, K33, K48, or K63). Two days post-transfection, cells were treated with 10 µM MG-132 for 3 hours at 37°C to inhibit proteasomal degradation. Myc-Ub levels on CFTR were detected using ELISA, as described above, with an anti-Myc antibody (9E10, Wako). CFTR ubiquitination levels were then normalized to total CFTR levels, which were measured using an anti-HA antibody.

### Pull-down experiment

To detect the interaction between HBH-ΔF508-CFTR-3HA and AMFR or SYVN1, 293MSR cells stably expressing HBH-ΔF508-CFTR-3HA were transfected with 10 nM siNC or siAMFR-1. Four days post-transfection, cells were treated with 10 µM MG-132 for 3 hours at 37°C, then solubilized in mild lysis buffer (150 mM NaCl, 20 mM Tris, 0.1% NP-40, pH 8.0) supplemented with 1 mM PMSF, 5 µg/mL leupeptin, and 5 µg/mL pepstatin. To examine the interaction between AMFR and CFTR domains, 293MSR RNF5/185 DKO cells were transfected with MB-AMFR along with CFTR NBD1-HiBiT, MSD1-HiBiT, or MSD2-HiBiT. Two days post-transfection, cells were treated with 10 µM MG-132 for 3 hours at 37°C, then solubilized in mild lysis buffer as described above. The cell lysates were incubated with NeutrAvidin agarose beads (ThermoFisher) for 2 hours at 4°C. After four washes with mild lysis buffer, the protein complexes were eluted in urea elution buffer (8 M urea, 2% SDS, 3 mM biotin) at room temperature for 30 minutes and analyzed by Western blotting, as shown in the figures.

### HiBiT and Nluc degradation assay

The HiBiT and Nluc degradation assays were performed as previously described.^29,34^ Briefly, proteins fused with HiBiT tag in the cytoplasmic region (CT) and cytosolic LgBiT (pBiT1.1-N [TK/LgBiT], Promega) were transiently expressed in 293MSR and RNF5/185 DKO cells transfected with the siRNA indicated in the figure legends. Four days post-transfection, cells seeded in 96-well plates were incubated with 50 µL of 0.1x Nano-Glo^®^ Endurazine (Promega) in CO₂-independent medium for 2.5 hours at 37°C in 5% CO₂. For the Nluc degradation assay, subconfluent BEAS-2B cells stably expressing Dox-inducible ΔF508-CFTR-Nluc(CT) were transfected with siRNA, and the expression of ΔF508-CFTR-Nluc(CT) was induced by treating the cells with 1 µg/mL Dox for 2 days. To measure the degradation kinetics, 10 µL of 600 µg/mL cycloheximide (CHX) was added to each well, resulting in a final concentration of 100 µg/mL. Luminescence was measured continuously at 5-minute intervals for 3-12 hours using a Luminoskan plate reader (ThermoFisher).

For the inhibitor assay, BTZ (100 nM) or BafA1 (1 µM) was pre-treated with Nano-Glo^®^ Endurazine (Promega) for 2.5 hours and was also added during the CHX chase. The luminescence signal of CHX-treated cells was normalized to the signal of untreated cells to calculate the remaining ERAD substrates during the CHX chase as described previously.^29^ The ERAD rate was calculated by fitting the luminescence decay to a one-phase exponential decay function using GraphPad Prism 8 (GraphPad Software, La Jolla, CA, USA).

### HiBiT retrotranslocation and ER disappearance assay

The retro-translocation and ER disappearance assays were performed as previously described.^29^ Briefly, proteins fused with HiBiT in the ER-luminal region and cytosolic LgBiT (retrotranslocation assay) or ER-luminal LgBiT (CNXss-LgBiT-KDEL, ER disappearance assay) were transiently expressed in 293MSR and RNF5/185 DKO cells transfected with the siRNA indicated in the figure legends. Four days post-transfection, cells seeded in 96-well plates were incubated with 50 µL of 0.5x Nano-Glo^®^ Endurazine (Promega) for the retrotranslocation assay or 0.1x Nano-Glo^®^ Endurazine for the ER disappearance assay, in CO₂-independent medium for 2.5 hours at 37°C in 5% CO₂. To detect retrotranslocation, 10 µL of 60 µM MG-132 (final concentration 10 µM) in CO₂-independent medium was added to each well. Luminescence was measured continuously at 5-minute intervals for 3 hours using a Luminoskan plate reader (ThermoFisher). The amount of retrotranslocated CFTR was calculated by subtracting the luminescence signal before MG-132 treatment from the signal after MG-132 treatment. The retrotranslocation rate was determined by linear fitting of the increase in retrotranslocation during MG-132 treatment. For the ER disappearance assay of CFTR-HiBiT(Ex), 10 µL of 600 µg/mL CHX in CO₂-independent medium was added to each well, reaching a final concentration of 100 µg/mL. Luminescence was measured continuously at 5-minute intervals for 3 hours using the Luminoskan plate reader (ThermoFisher). The luminescence signal from CHX-treated cells was normalized to the signal from untreated cells as described previously.^29^ The ER disappearance rate was calculated by fitting the luminescence decay to a one-phase exponential decay function using GraphPad Prism 8.

### Subcellular fractionation

Microsomes were isolated from 293MSR WT and RNF5/185 DKO cells stably expressing HBH-ΔF508-CFTR-3HA. The cells were resuspended in resuspension buffer (10 mM HEPES, pH 7.5, 0.25 M sucrose, 10 mM KCl, 1.5 mM MgCl₂, 1 mM EDTA, 1 mM EGTA, 5 µg/mL leupeptin, 5 µg/mL pepstatin A, and 1 mM PMSF) and sheared by passing the suspension 30 times through a 25G needle. The cell homogenates were then centrifuged at 1,000 g at 4°C for 5 minutes. The supernatant was recentrifuged at 100,000 g at 4°C for 1 hour. The resulting supernatant was used as the cytosol, and the pellet fraction containing microsomes was dissolved in RIPA buffer. Both microsomal and cytosolic fractions were analyzed by Western blotting.

### Statistical analysis

Data from at least three independent experiments were used for quantification, with results presented as mean ± standard deviation (SD). Statistical significance was determined from at least three biological replicates (n) using one-way or two-way ANOVA, or a two-tailed Student’s t-test, as indicated in the figure legends, and analyzed with GraphPad Prism 8. A P-value < 0.05 was considered statistically significant. While data distribution was assumed to be normal, this was not formally tested.

---

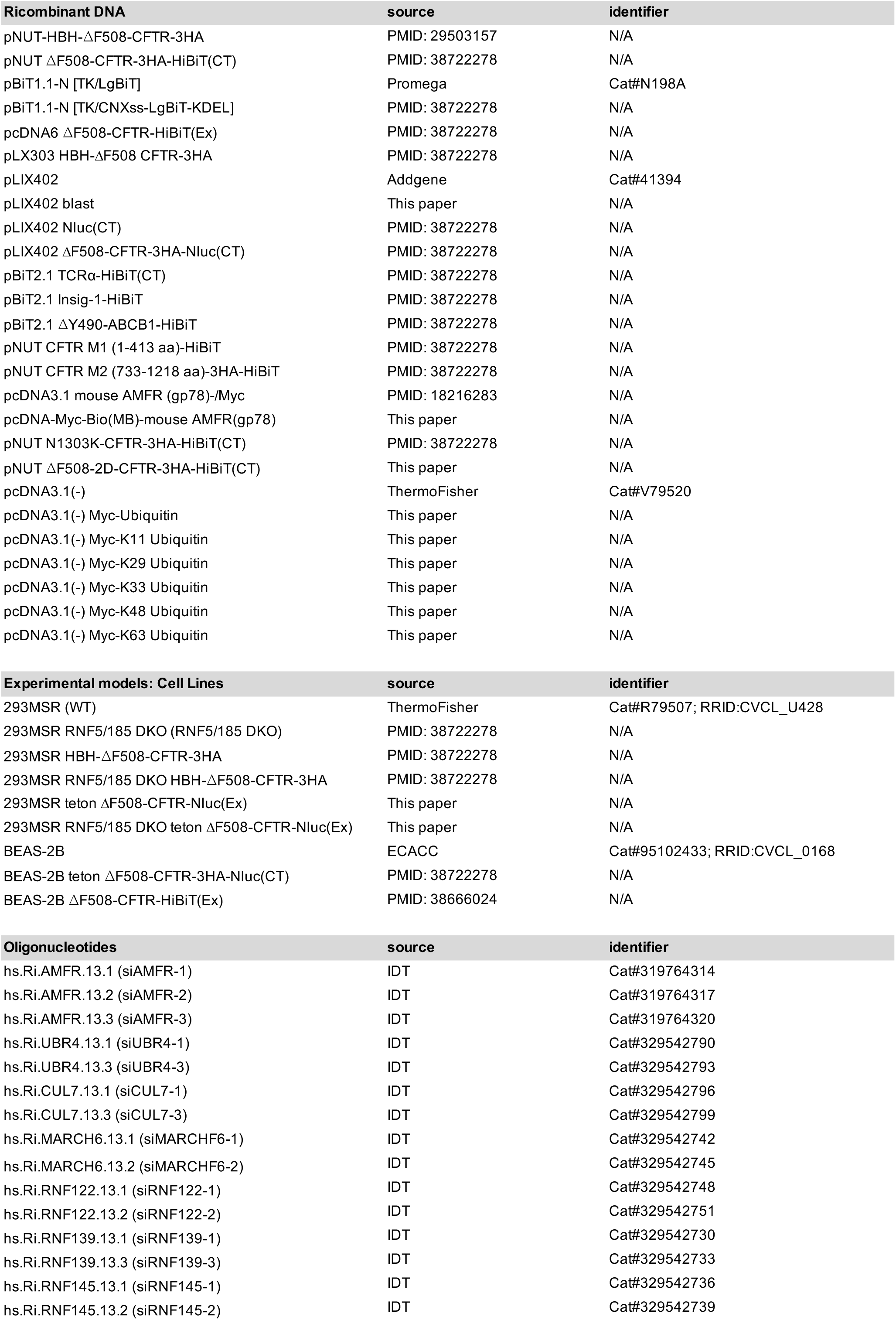
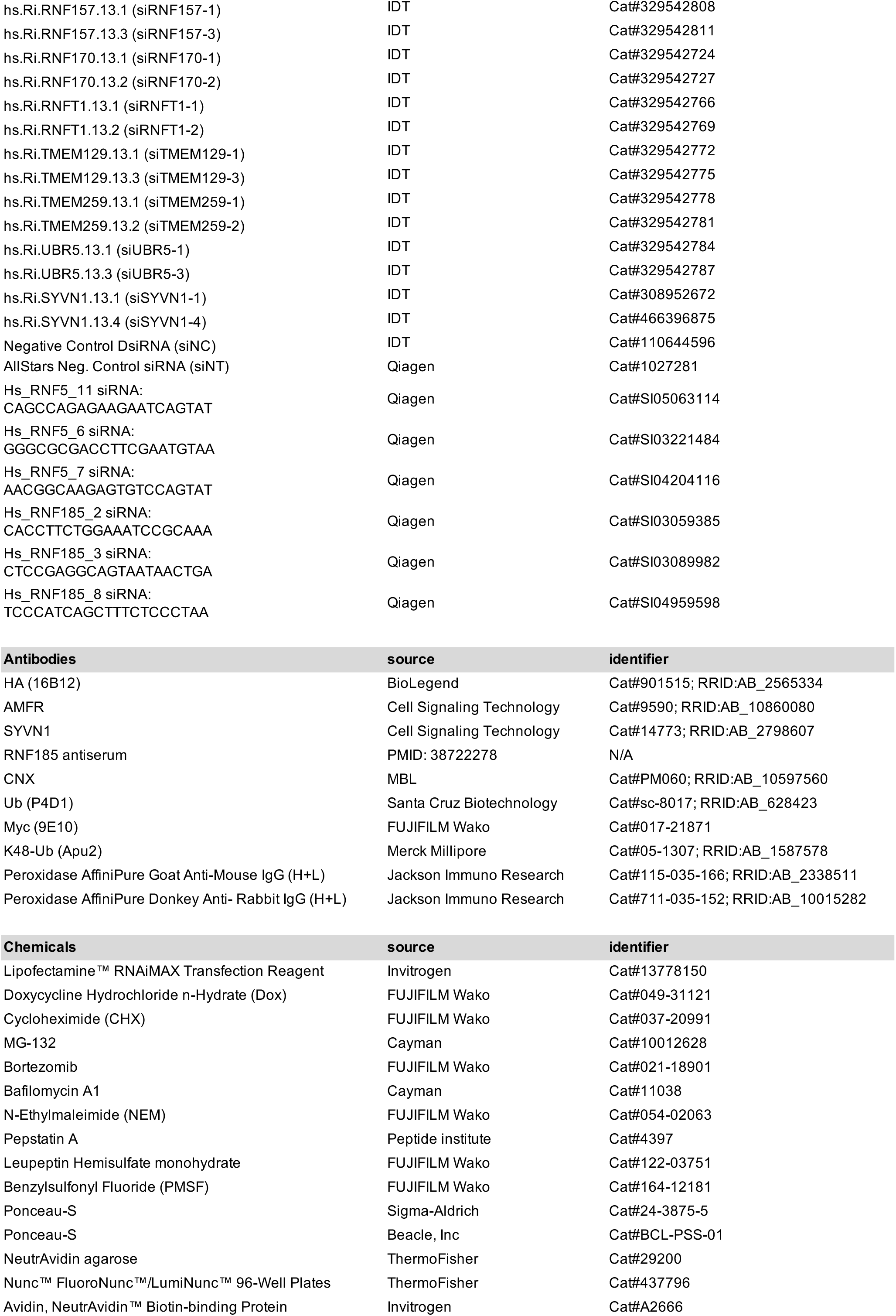
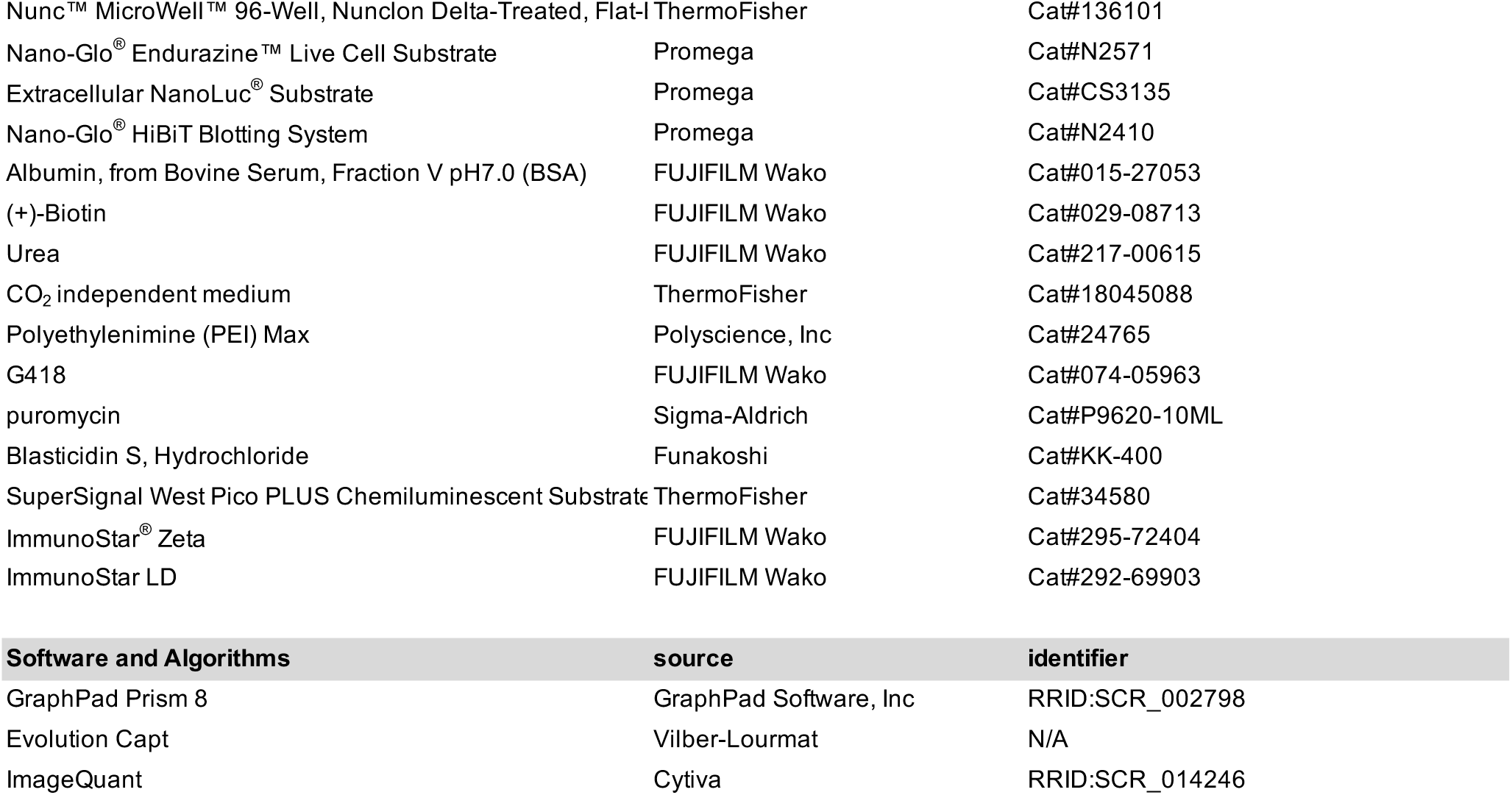

