## supplemental information for "AMFR provides an ERAD bypass mechanism to maintain proteostasis under canonical E3 ligase deficiency"

#### Supplemental methods

##### Structural representation

The structural models of the CFTR–RNF5–RNF185 and CFTR–AMFR complexes were generated using the AlphaFold3 program (https://alphafoldserver.com). The predictions were based on the amino acid sequences of CFTR (NP_000483.3), RNF5 (NP_008844.1), RNF185 (NP_689480.2), and AMFR (NP_001135.3). All structural representations were prepared using PyMOL (The PyMOL Molecular Graphics System, Version 3.0.0, Schrödinger, LLC). The amino acid residues in the CFTR TM region that could interact with RNF5, RNF185, or AMFR were defined as those located within 5 Å of the respective ligase in the predicted complex structures.

#### Supplemental Figure legends

##### Figure S1.

**Effects of selected E3 ligase KD on cell surface ΔF508-CFTR-Nluc(Ex) in 293MSR WT cells**

The assay was performed as described in Fig. 1B, following transfection with 10 nM siNC (negative control) or pooled siRNAs (n=3). Statistical significance was assessed using one-way ANOVA with Dunnett's multiple comparisons test. Data are presented as mean ± SD. ns, not significant.

##### Figure S2.

**ER stress markers increased in RNF5/185 DKO cells**

Expression levels of ER stress markers were analyzed by RNA-seq. mRNA levels of ER stress-related genes in 293MSR WT and RNF5/185 DKO cells were quantified as shown in Figure 1A, with transcript abundance represented as the TPM ratio (DKO/WT) (n = 1).

##### Figure S3.

**Predicted structures of CFTR–RNF5–RNF185 and CFTR–AMFR complexes**

(A, B) Structural models of the CFTR–RNF5–RNF185 and CFTR–AMFR complexes were predicted using AlphaFold (AF). The transmembrane domain 1 (TM1) region of CFTR, potentially interacting with RNF5 or RNF185 (A), and AMFR (B), is highlighted in magenta and red, respectively.

(C) Amino acid residues in the CFTR TM1 region predicted to interact with RNF5 or RNF185 (top) and AMFR (bottom) are listed based on five individual AF models of each complex.

### Figure S1

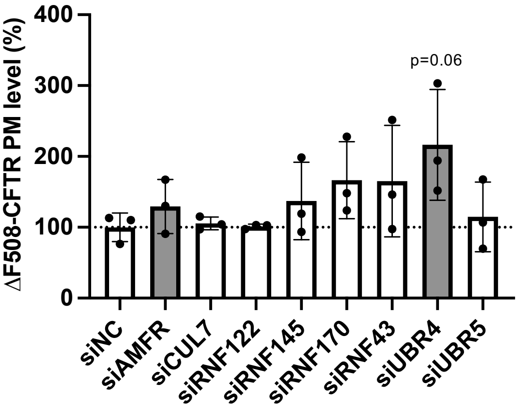
ΔF508-CFTR PM level (%)
400
300
200
100
0
p=0.06
siNC
siAMFR
siCUL7
siRNF122
siRNF145
siRNF170
siRNF43
siUBR4
siUBR5

**Figure S2**

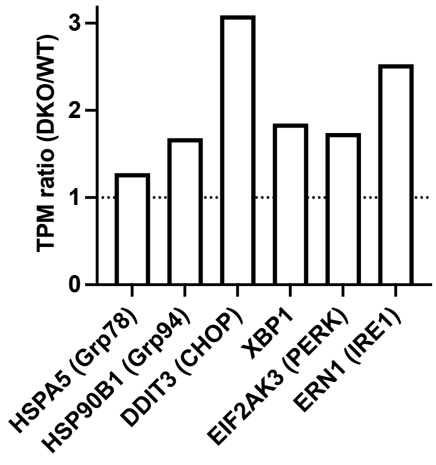

TPM ratio (DKO/WT)
0
1
2
3
HSPA5 (Grp78)
HSP90B1 (Grp94)
DDIT3 (CHOP)
XBP1
EIF2AK3 (PERK)
ERN1 (IRE1)

### Figure S3

A

**CFTR-RNF5-RNF185 complex AF model 0**

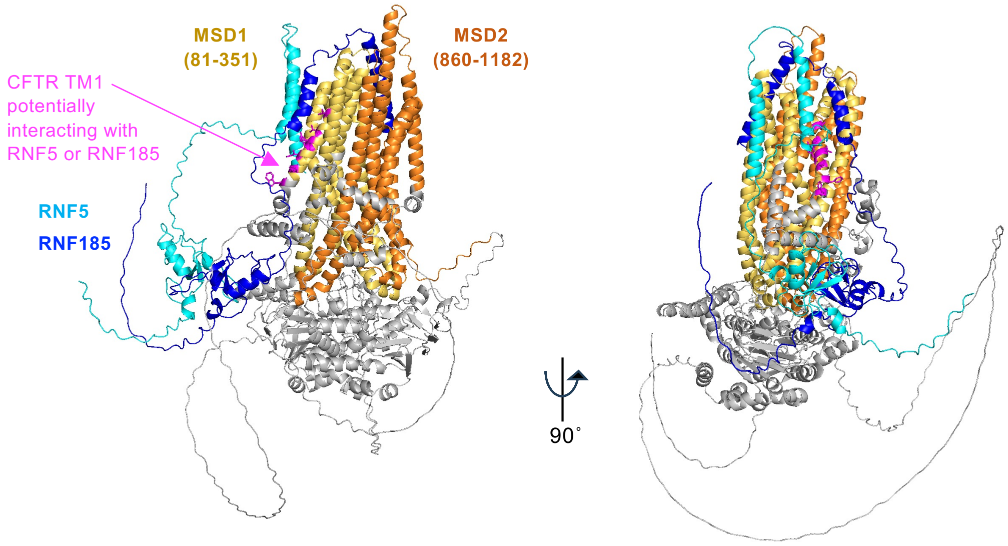


B

**CFTR-AMFR complex AF model 0**

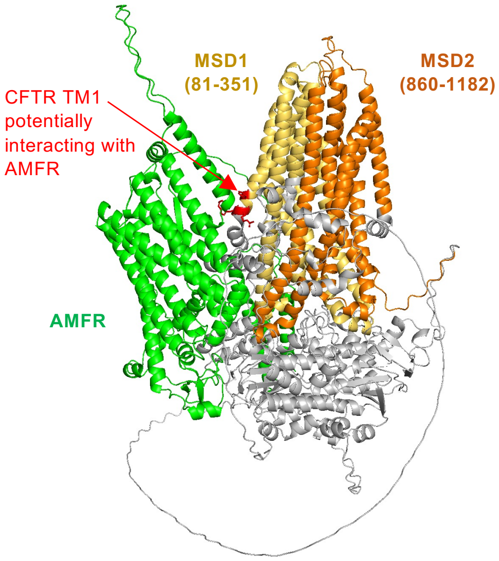


C

**CFTR-RNF5-RNF185 complex**

| | *AF model* | *CFTR TM1 (78-98 aa)* |
|---|---|---|
| **RNF5** | 0 | **FWRFMFYGIFLYLGEVTKAVQ** |
| | 1 | no binding |
| | 2 | no binding |
| | 3 | no binding |
| | 4 | **FWRFMFYGIFLYLGEVTKAVQ** |
| | *AF model* | *CFTR TM1 (78-98 aa)* |
| **RNF185** | 0 | **FWRFMFYGIFLYLGEVTKAVQ** |
| | 1 | no binding |
| | 2 | **FWRFMFYGIFLYLGEVTKAVQ** |
| | 3 | no binding |
| | 4 | **FWRFMFYGIFLYLGEVTKAVQ** |

**CFTR-AMFR complex**

| | *AF model* | *CFTR TM1 (78-98 aa)* |
|---|---|---|
| **AMFR** | 0 | **FWRFMFYGIFLYLGEVTKAVQ** |
| | 1 | **FWRFMFYGIFLYLGEVTKAVQ** |
| | 2 | **FWRFMFYGIFLYLGEVTKAVQ** |
| | 3 | **FWRFMFYGIFLYLGEVTKAVQ** |
| | 4 | **FWRFMFYGIFLYLGEVTKAVQ** |
